# PRMT5 as a Novel Druggable Vulnerability for EWSR1-ATF1-driven Clear Cell Sarcoma

**DOI:** 10.1101/2022.03.23.485409

**Authors:** Bingbing X. Li, Larry L. David, Lara E. Davis, Xiangshu Xiao

**Author notes:** **Disclosure:** B. X. L. and X. X. are named inventors on a US Provisional patent application on the use of PRMT5 inhibitors (# 63/208,655). All the other authors have nothing to disclose.

## Abstract

Clear cell sarcoma of soft tissue (CCSST) is an ultra-rare sarcoma with poor prognosis presently with no cure. It is characterized by a balanced t(12;22) (q13;q12) chromosomal translocation, resulting in a fusion of the Ewing’s sarcoma gene *EWSR1* with activating transcription factor 1 (*ATF1*) to give an oncogene *EWSR1-ATF1*. Unlike normal ATF1, whose transcription activity is dependent on phosphorylation, EWSR1-ATF1 is constitutively active to drive ATF1-dependent gene transcription to cause tumorigenesis. No EWSR1-ATF1-targeted therapies have been identified due to the challenges in targeting intracellular transcription factors. To identify potential druggable targets for CCSST, we show that protein arginine methyltransferase 5 (PRMT5) is a novel enzyme in enhancing EWSR1-ATF1-mediated gene transcription to sustain CCSST cell proliferation. Genetic silencing of *PRMT5* in CCSST cells resulted in severely impaired cell proliferation and EWSR1-ATF1-driven transcription. Furthermore, the clinical-stage PRMT5 inhibitor **JNJ-64619178** potently and efficaciously inhibited CCSST cell growth *in vitro* and *in vivo*. These results provide new insights into PRMT5 as a transcription regulator and warrant **JNJ-64619178** for further clinical development to treat CCSST patients.

## Introduction

Clear cell sarcoma of soft tissue (CCSST), first described in 1965,^1^ is an ultra-rare but aggressive soft tissue sarcoma that most often develops in the lower extremity close to tendons and aponeuroses of adolescents and young adults.^1-3^ The standard treatment for localized disease is to perform wide surgical resection or whole limb amputation. However, up to 85% of the patients develop the recurrence and eventual metastasis.^1, 4^ The 10-year overall survival rate is only ∼35% and the 5-year survival rate is only ∼20% for metastatic disease.^5-8^ Since most of the patients are adolescents and young adults, the overall life years lost due to CCSST are more substantial. CCSST is notorious for its insensitivity to existing chemotherapies or radiation therapy. No targeted therapies exist for CCSST.

The hallmark of CCSST is characterized by a balanced t(12;22) (q13;q12) chromosomal translocation, which results in a gene fusion between the Ewing’s sarcoma gene *EWSR1* and activating transcription factor 1 (*ATF1*) to give an oncogene *EWSR1-ATF1*.^9-11^ EWSR1 is known as an RNA-binding protein.^12^ ATF1 is a member of the cAMP-responsive element (CRE) binding protein (CREB) family transcription factor.^13, 14^ The gene fusion retains the C-terminal basic leucine-zipper (bZIP) DNA-binding domain of ATF1. The CREB family of transcription factors requires phosphorylation at Ser133 of CREB1 or Ser63 of ATF1 by protein Ser/Thr kinases including cAMP-regulated protein kinase A (PKA) in order to be transcriptionally active.^15^ However, in the case of EWSR1-ATF1 fusion, the transcription activation domain present in EWSR1 renders it constitutively active in driving CREB1/ATF1-depenent gene transcription through its bZIP domain independent of phosphorylation.^16^

Genetic mouse modeling studies have established that expression of EWSR1-ATF1 is sufficient to drive the development of CCSST in mice, resembling human pathologies.^17-19^ Besides the initial discovery of type 1 *EWSR1-ATF1* fusion in CCSST patients, 6 other types of *EWSR1-ATF1* fusion and *EWSR1-CREB1* have also been reported in CCSST patients.^20^ Furthermore, these gene fusions have been observed in other CCS-like disease pathologies in other parts of the body including gastrointestinal tract and other abdominal surfaces.^4^ Together, the identification of *EWSR1-ATF1* and *EWSR1-CREB1* fusions underscores the critical role of constitutive CREB1/ATF1-mediated gene transcription in the development of CCSST and other CCS-like cancers that lack effective therapies. However, it is unclear how EWSR1-ATF1 can activate CREB1/ATF1-dependent gene transcription without phosphorylation. Lack of this mechanistic understanding inhibits our capability to identify novel therapeutics for these sarcomas. In this paper, we describe the identification of protein arginine methyltransferase 5 (PRMT5) as a new EWSR1-ATF1 binding co-activator to stimulate its transcription activity. We further employed genetic and chemical means to inhibit PRMT5 to demonstrate that PRMT5 is a novel druggable vulnerability in EWSR1-ATF1-driven CCSST.

## Results

### EWSR1-ATF1 is intrinsically disordered but constitutively active in driving CREB1/ATF1-mediated gene transcription

The most common fusion between EWSR1 and ATF1 generates EWSR1(2-325)-ATF1(66-271) fusion containing N-terminal portion of EWSR1 and C-terminal portion of ATF1 with its intact bZIP DNA-binding domain.^21^ Genetic studies have provided compelling evidence that the growth and survival of CCSST cells are critically dependent on the continuous expression of EWSR1-ATF1 and its transcription activity.^17, 22^ As a fusion protein between an RNA-binding protein EWSR1 and a transcription factor ATF1, the major portion of EWSR1-ATF1 fusion protein is predicted to be intrinsically disordered. Indeed, when the fusion protein sequence was analyzed by PONDR (Predictor of Natural Disordered Regions) algorithm,^23^ the majority of the EWSR1 region is found to be disordered (Figure 1A). As expected, the bZIP domain in the C-terminal portion of ATF1 is ordered. Consistent with this prediction, the artificial intelligent (AI) algorithm AlphaFold^24^ also predicts that EWSR1 portion is unstructured while a good fraction of ATF1 including the bZIP domain is structured (Figure 1B). Even within the structured region of ATF1, no obvious small molecule binding sites can be readily identified (Figure S1A). As a consequence, direct targeting of EWSR1-ATF1 by small molecules is challenging and no direct small molecule inhibitors of EWSR1-ATF1 have been developed.

**Figure 1.**
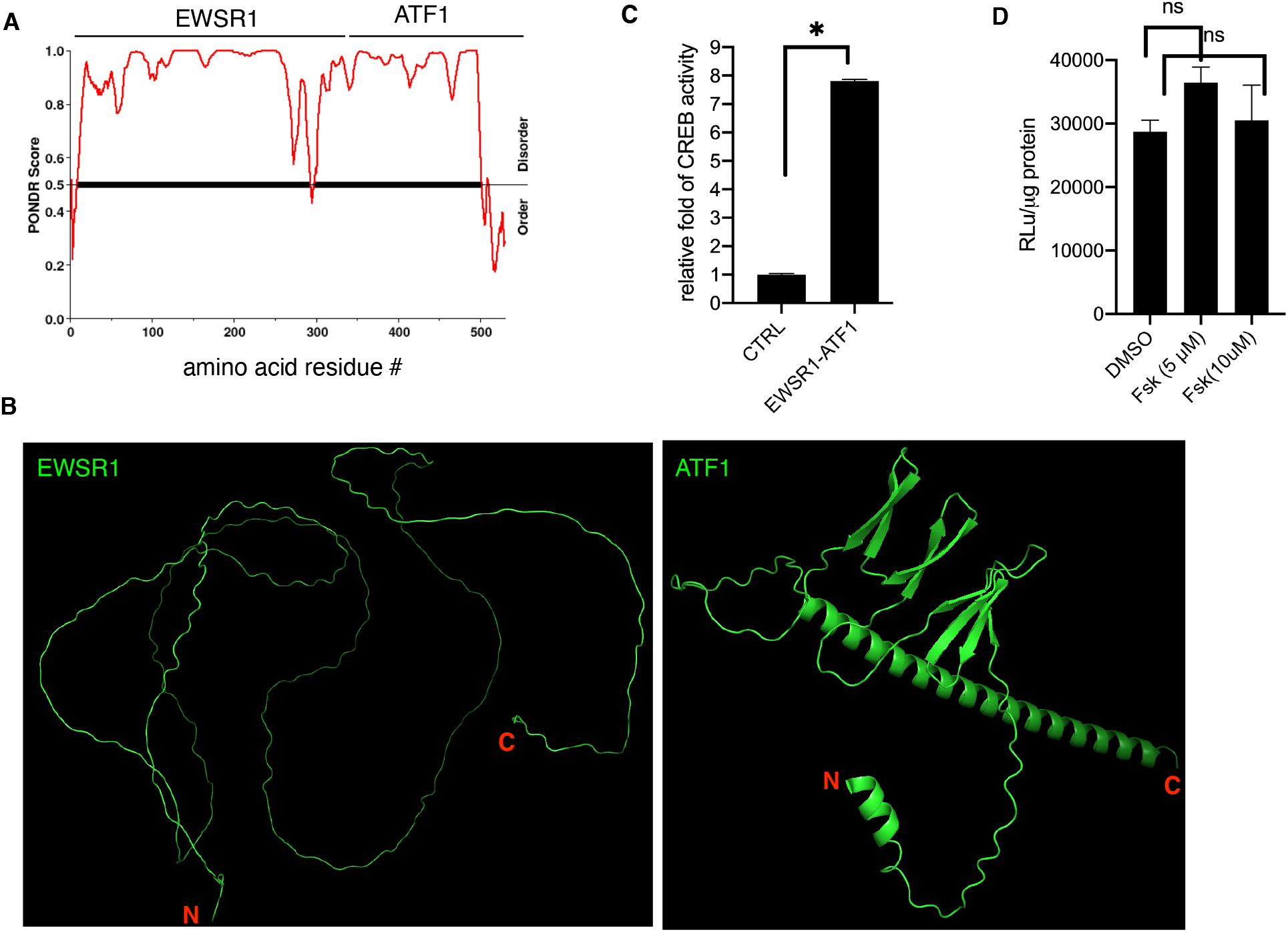
EWSR1-ATF1 is intrinsically disordered and constitutively active in driving CREB1/ATF1-dependent gene transcription. (A) EWSR1-ATF1 is an intrinsically disordered protein. The PONDR score was calculated using online PONDR server at http://www.pondr.com. (B) Cartoon presentation of AlphaFold predicted structures of EWSR1(2-325) (left) and ATF1(66-271) (right). The N- and C-termini are labeled. (C) EWSR1-ATF1 is constitutively active. HEK 293T cells were transfected with an empty vector or EWSR1-ATF1 along with CREB1 transcription reporter plasmid CRE-RLuc. Then the renilla luciferase activity was measured and normalized to the protein content. (D) EWSR1-ATF1’s transcription activity is independent of phosphorylation. HEK 293T cells were transfected with EWSR1-ATF1 along with CREB1 transcription reporter plasmid CRE-RLuc. The cells were treated with different concentration of forskolin (Fsk) for 5 h before the renilla luciferase activity was measured, which was further normalized to the protein content. * *P* < 0.05.

Despite EWSR1-ATF1 is largely disordered, it has been reported that it functions as a constitutively active transcription factor.^16^ To confirm this, we created the in-frame Flag-tagged fusion between EWSR1(2-325) and ATF1(66-271) (Figure S1B). When EWSR1-ATF1 was expressed along with our previously described CREB1/ATF1 transcription reporter plasmid (CRE-RLuc) in HEK 293T cells,^25^ we found that the exogenously expressed EWSR1-ATF1 is transcriptionally active in driving CREB1/ATF1-depedent gene transcription (Figure 1C). Furthermore, this transcription activity is independent of phosphorylation as adding PKA activator forskolin (Fsk) did not further enhance its transcription activity (Figure 1D). These results support that EWSR1-ATF1 is constitutively active in driving CREB1/ATF1-dependent gene transcription.

### EWSR1-ATF1 interacts with PRMT5

Since EWSR1-ATF1 has been established as the genetic driver for CCSST development and maintenance, targeting this driver represents a potentially promising approach to develop effective therapeutics. However, the intrinsically disordered nature of EWSR1-ATF1 has created challenges in developing direct small molecule inhibitors to inhibit its transcription activity as potential therapeutics. To circumvent this intrinsic challenge, we sought to identify critical proteins that would interact with EWSR1-ATF1 to support its constitutive transcription activity and hypothesized that these proteins might serve as potential drug targets for CCSST. Therefore, we expressed Flag-tagged EWSR1-ATF1 in HEK 293T cells. To identify the proteins that would interact with EWSR1-ATF1, the expressed EWSR1-ATF1 was subjected to immunoprecipitation by anti-Flag (M2) followed by mass spectrometry (IP-MS) analysis (Figure 2A). An IgG control pulldown was also included. This analysis identified 157 proteins that presented higher abundance in the anti-Flag sample (Table S1). Gene ontology (GO) analysis of these 157 proteins in the biological processes using g:Profiler^26^ revealed that many processes involving RNA molecules are enriched (Table S2). The top 15 biological processes enriched are shown in Figure 2B, which include both RNA catabolic and metabolic processes. Among these processes, the mRNA metabolic process contains protein arginine methyltransferase 5 (PRMT5) that has been recently become a druggable target.^27^ Various PRMT5 inhibitors have been developed and at least 5 of them are in Phase 1/2 clinical trials for different malignancies.^28^ Additionally, PRMT5 is one of only 6 proteins (PPM1B, KIF11, PRMT5, ACTR3, HBB, PTGES3) that interact with EWSR1-ATF1 specifically and have 3 or more unique peptide identifications (Figure 2C, Figure S2 and Table S1). Furthermore, while PRMT5 has not previously been shown to interact with EWSR1-ATF1, it was reported that PRMT5 could function as a transcription co-activator for CREB1 in hepatocytes to enhance CREB1-mediated gene transcription.^29^ For these reasons, we focused our investigation into PRMT5 as a putative critical target for EWSR1-ATF1-driven CCSST.

**Figure 2.**
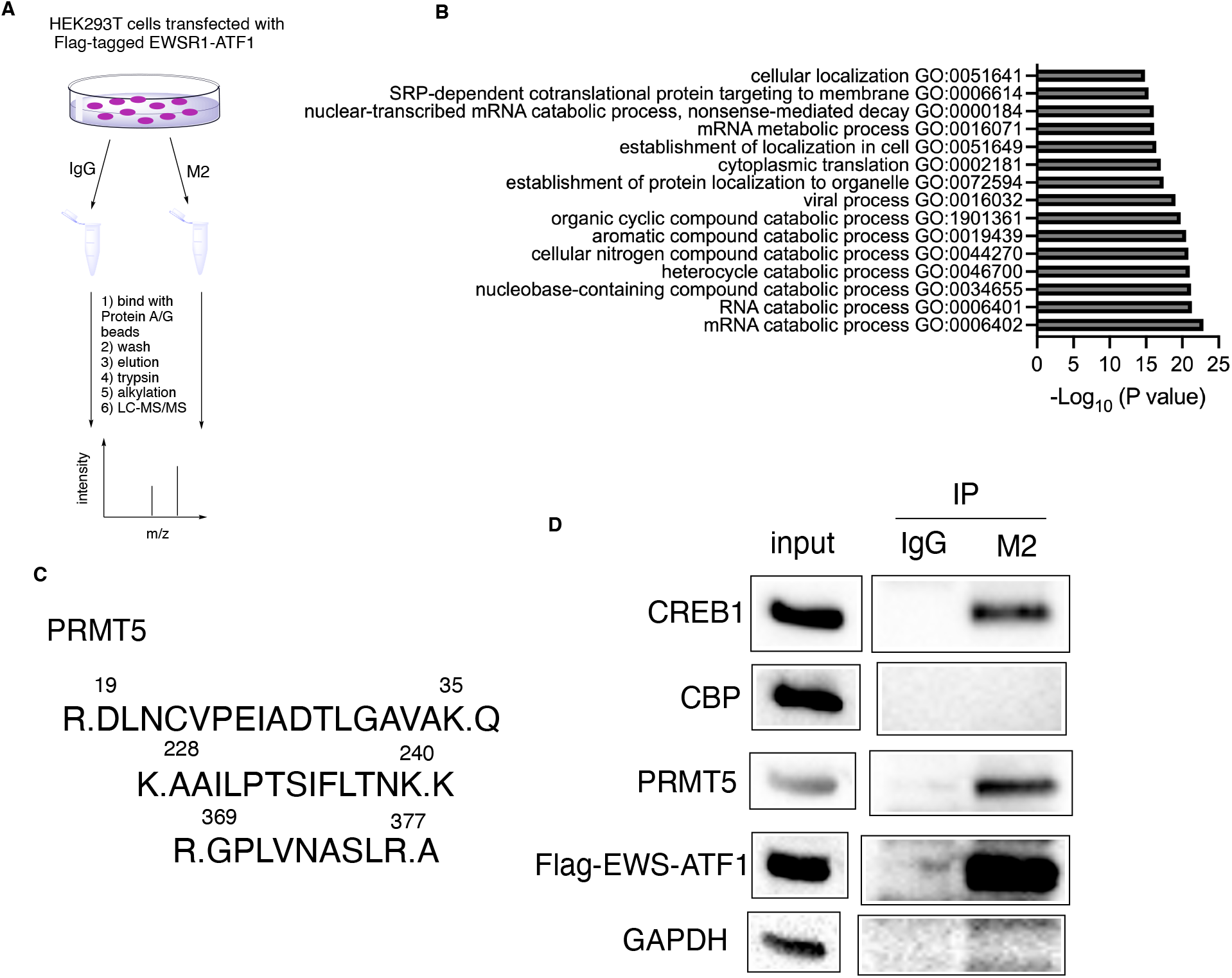
EWSR1-ATF1 interacts with PRMT5. (A) A schematic diagram of IP-MS/MS to identify proteins that interact with EWSR1-ATF1. (B) Top 15 enriched biological processes by GO analysis of the 167 proteins interacting with EWSR1-ATF1 using g:Profiler. The enriched biological processes were rank ordered by adjusted *P* values. See Table S2 for the full list. (C) Identification of PRMT5 to interact with EWSR1-ATF1. HEK 293T cells were transfected with Flag-tagged EWSR1-ATF1. Then the lysates were prepared for co-immunoprecipitation (co-IP) using either IgG or anti-Flag (M2). The eluted proteins were analyzed by LC-MS/MS. The identified peptides corresponding to PRMT5 are shown (from M2 IP). No peptides corresponding to PRMT5 were identified in the IgG immune complex. (D) Validation of the interaction between EWSR1-ATF1 and PRMT5. Flag-tagged EWS-ATF1 was expressed in HEK 293T cells and the lysates were processed in the same way as in (A). The bound proteins were analyzed by Western blot using indicated antibodies.

PRMT5 is a type II protein arginine methyltransferase leading to symmetric arginine dimethylation of both histone and non-histone proteins to regulate a diverse range of activities including gene transcription, RNA splicing and DNA repair.^30^ To confirm the proteomics findings, we expressed Flag-tagged EWSR1-ATF1 in HEK 293T cells and immunoprecipitated the cell lysates using anti-Flag antibody (M2). Then the bound proteins were analyzed by Western blot. Indeed, strong PRMT5 signal was detected in the M2 immunoprecipitate, but not in the control IgG immunoprecipitate (Figure 2D). CREB1 can form a heterodimer with the C-terminal domain of ATF1 through its bZIP domain that is retained in EWSR1-ATF1. Indeed, CREB1 was detected in the M2 immune complex (Figure 2D). Conflicting reports exist to show that EWSR1-ATF1 constitutively interacts with CREB-binding protein (CBP) to support its constitutive transcription activity.^31, 32^ In our co-IP analysis, the constitutive interaction between EWSR1-ATF1 and CBP was not detected. As a control, GAPDH was not detected in either of the immune complexes (Figure 2B). Together, these results demonstrate that EWSR1-ATF1 interacts with PRMT5 in the cells to possibly function as a transcription co-activator to support EWSR1-ATF1-mediated gene transcription.

### PRMT5 is critical for EWSR1-ATF1-mediated gene transcription

The results presented above suggest that PRMT5 might be a potential transcription co-activator for EWSR1-ATF1. To test if PRMT5 is bound to the gene promoter region of EWSR1-ATF1 target gene *c-Fos*^22^ in CCSST cells, chromatin immunoprecipitation (ChIP) was performed in CCSST cell line DTC-1. After the cells were crosslinked with formaldehyde, they were subjected to ChIP analysis using IgG, anti-CREB1 or anti-PRMT5. CREB1 forms a heterodimer with EWSR1-ATF1 (Figure 2D) and served as a positive control for the ChIP assay. Indeed, the CRE site in the *c-Fos* promoter region is occupied by CREB1 (Figure 3A). Importantly, this site is also occupied by PRMT5 (Figure 3A), suggesting that PRMT5 may enhance EWSR1-ATF1’s transcription activity by binding to the promoter region of its target genes.

**Figure 3.**
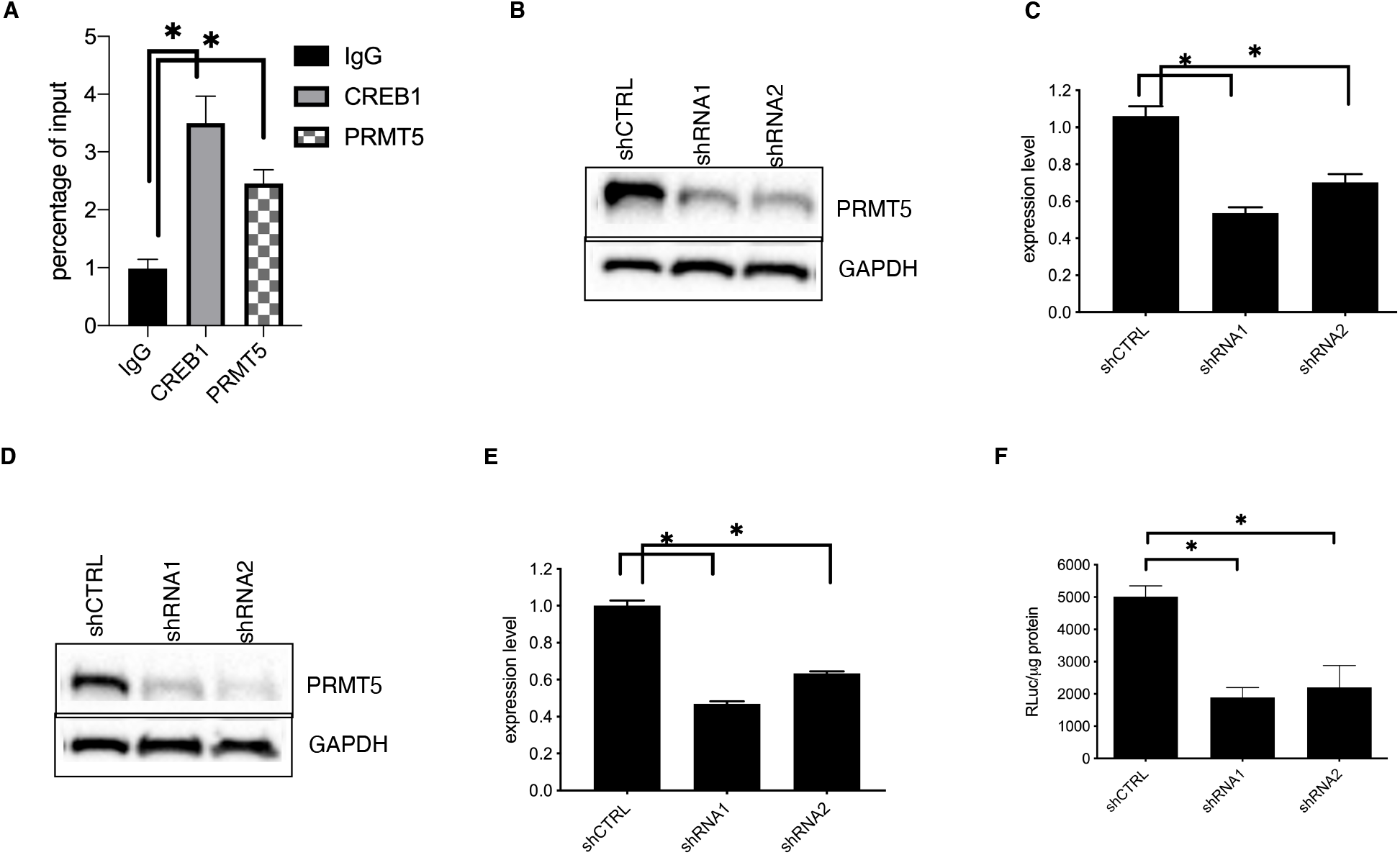
PRMT5 is critical for EWSR1-ATF1-mediated gene transcription. (A) PRMT5 is bound to the *c-Fos* promoter in DTC-1 cells. DTC-1 cells were subjected to ChIP assay protocol as described in the Methods. The enriched DNA by individual antibodies was analyzed by qPCR analysis. (B) The expression of *PRMT5* was silenced using two different shRNA constructs in SU-CCS-1 cells. (C) The expression of *c-Fos* was reduced with sh*PRMT5*. SU-CCS-1 cells were transduced with indicated lentiviruses expressing different shRNA constructs. The total RNA was collected for qRT-PCR analysis. (D) Silencing *PRMT5* expression in DTC-1 cells using shRNA. DTC-1 cells were transduced with lentiviruses expressing indicated shRNA constructs. The lysates were prepared for Western blot analysis using indicated antibodies. (E) Silencing *PRMT5* expression in DTC-1 cells decreased mRNA level of *c-Fos*. The cells transduced with shRNA lentiviruses were processed for qRT-PCR analysis for quantification of *c-Fos* transcript. (F) EWSR1-ATF1-driven CRE-RLuc expression is reduced by sh*PRMT5*. DTC-1 cells were transduced with lentiviruses expressing CRE-RLuc along with lentiviruses expressing indicated shRNA. Then the renilla luciferase activity was quantified and normalized to protein concentration under each condition. * *P* < 0.05.

To investigate if PRMT5 is required for sustained transcription of *c-Fos* in CCSST cells, its expression was silenced in SU-CCS-1 cells using two different shRNA constructs. Both of these constructs efficiently knocked down the expression of PRMT5 (Figure 3B). When the expression of PRMT5 was reduced, the expression of EWSR1-ATF1’s target gene *c-Fos* was also significantly reduced, as assessed by quantitative reverse transcription-polymerase chain reaction (qRT-PCR) analysis (Figure 3C), suggesting that PRMT5 is critical for enhancing EWSR1-ATF1’s transcription activity. Similar knocking down results were also obtained in another CCSST cell line DTC-1 (Figure 3D-E). We further evaluated PRMT5’s role in supporting in EWSR1-ATF1-mediated gene transcription using a CREB1/ATF1 transcription reporter assay in DTC-1 cells. The cells were transduced with a lentivirus containing renilla luciferase reporter gene under the control of three tandem copies of CRE that could bind EWSR1-ATF1.^25^ Upon *PRMT5* knockdown by two different shRNAs, the CREB1/ATF1 transcription reporter activity was significantly reduced (Figure 3F). These evaluations of the transcription reporter and endogenous gene expression in CCSST cells showed that PRMT5 is critical for the constitutive EWSR1-ATF1-driven transcription.

### PRMT5 inhibition leads to CCSST cell growth inhibition

Because EWSR1-ATF1-driven transcription is critical for sustained CCSST cell proliferation, we investigated if silencing the expression of *PRMT5* would impact CCSST cellular proliferation. To this end, the expression of *PRMT5* was silenced in both SU-CCS-1 and DTC-1 cells using two independent shRNA constructs. Upon *PRMT5* knockdown (Figure 3B and 3D), the proliferation of both SU-CCS-1 and DTC-1 cells was significantly impaired (Figure 4A), suggesting that PRMT5 is critical for the proliferation of CCSST cells.

**Figure 4.**
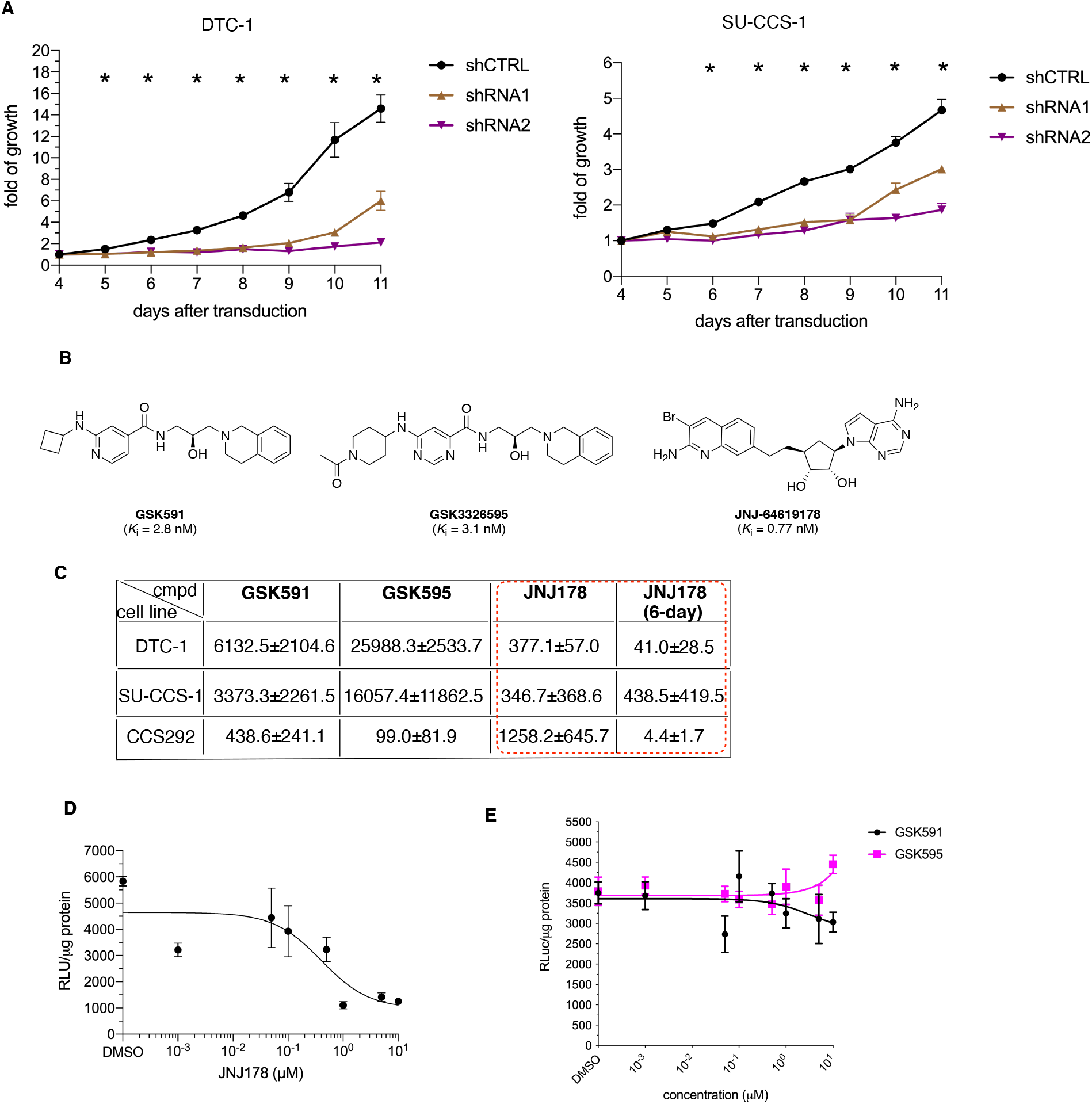
Inhibition of PRMT5 leads to impaired CCSST cell growth. (A) Silencing *PRMT5* expression resulted in reduced cellular proliferation in DTC-1 (left) and SU-CCS-1 (right) cells. The cells were transduced with indicated shRNA constructs. After 3 days, the cell growth was analyzed by the MTT reagent over a period of 8 days. (B) Chemical structures of PRMT5 inhibitors to be used in this paper. Their apparent inhibition constants *K*_i_ are shown and from published references. (C) Summary of the GI_50_s (nM) of 3 different PRMT5 inhibitors in 3 different CCSST cell lines. The cells were treated with different concentrations of the indicated compounds for either 3 days or 6 days (as indicated). Then the remaining viable cells were quantified by the MTT reagent. The values represent the mean + SD of at least 2 independent experiments performed in triplicates. See Figure S4 for their respective dose-response curves. (D) **JNJ-64619178** inhibited EWSR1-ATF1-mediated gene transcription in DTC-1 cells. DCT-1 cells with CRE-RLuc reporter integration were treated with different concentrations of **JNJ-64619178** for 24 h. Then the remaining renilla luciferase activity was measured and normalized to protein content in each well. (E) **GSK591** and **GSK3326595** did not inhibit EWSR1-ATF1-mediated gene transcription in DTC-1. DTC-1 cells with CRE-RLuc were treated with different concentrations of GSK compounds for 24 h. Then the remaining renilla luciferase activity was measured and normalized for protein content in each well. * *P* < 0.05.

Given the potentially important role of PRMT5 in CCSST cells and druggability of PRMT5,^27^ we investigated the potential efficacy of PRMT5 inhibitors in inhibiting CCSST cell growth. We selected 3 highly potent and selective PRMT5 inhibitors representing two different modes of inhibition of PRMT5 to assess their efficacy in CCSST models. **GSK591** and **GSK3326595** bind at the protein substrate binding pocket^33, 34^ while **JNJ-64619178** binds simultaneously to the *S*-adenosyl methionine (SAM)- and protein substrate-binding pockets (Figure 4B and Figure S3).^35^ The inhibitors were tested in 3 different CCSST cell lines: SU-CCS-1, DTC-1 and CCS292. The cells were incubated with different PRMT5 inhibitors for either 3 days or 6 days. Then the viable cells were quantified using the MTT reagent and the GI_50_ values were calculated, which are the concentrations needed to inhibit the cell growth by 50%.^36^ As shown in Figure 4C and Figure S4A-C, different PRMT5 inhibitors showed differential sensitivity profiles in the panel of CCSST cell lines. **GSK591** and **GSK3326595** were only weakly active in DTC-1 and SU-CCS-1 cells with GI_50_s in the high μM concentration range although its activity in CCS292 cells was much more potent. On the other hand, **JNJ-64619178** displayed more potent activities in both DTC-1 and SU-CCS-1 cells with GI_50_ = 377 and 347 nM, respectively, during a 3-day incubation period. Thus, **JNJ-64619178** was further evaluated in a 6-day treatment protocol. Upon this extended exposure, **JNJ-64619178** showed dramatically improved activity with GI_50_ = 41.0, 438.5, and 4.4 nM in DTC-1, SU-CCS-1 and CCS292 cells, respectively (Figure 4C and Figure S4D). While all the three inhibitors are highly potent in inhibiting the methyltransferase activity of PRMT5,^33-^ _35_ the differential sensitivity profile among the different PRMT5 inhibitors in CCSST cells suggest that the different binding modes of these inhibitors can uniquely modulate EWSR1-ATF1-mediated gene transcription.

**JNJ-64619178**’s potent growth inhibitory activity in CCSST cells prompted us to further test its capability to inhibit EWSR1-ATF1-mediated gene transcription. In the CREB1/ATF1 transcription reporter assay in DTC-1 cells, **JNJ-64619178** displayed an IC_50_ of 422 nM (Figure 4D). Consistent with the low potency of GSK compounds in inhibiting CCSST cell growth, neither of these two substrate-competitive inhibitors significantly inhibited EWSR1-ATF1’s transcription activity (Figure 4E), further emphasizing the unique mechanism of action of **JNJ-64619178** in CCSST cells.

Altogether, these results support a critical role of PRMT5 in enhancing EWSR1-ATF1’s transcription activity for sustained cell growth in CCSST cells and suggest that PRMT5 is a potential novel druggable vulnerability in CCSST cells.

### JNJ-64619178 inhibits EWSR1-ATF1-mediated gene transcription and CCSST tumor growth *in vivo*

The results presented above strongly suggest that **JNJ-64619178** may represent a novel therapeutic agent for the deadly CCSST. We first examined if **JNJ-64619178** could downregulate the expression of endogenous EWSR1-ATF1’s target gene in SU-CCS-1 cells. When the cells were treated with **JNJ-64619178**, we observed that **JNJ-64619178** dose-dependently inhibited transcription of *c-Fos* (Figure 5A). **JNJ-64619178** treatment in DTC-1 cells caused a similar decrease in the mRNA level of *c-Fos* (Figure 5B). Decreased transcription of *c-Fos* by **JNJ-64619178** in SU-CCS-1 and DTC-1 cells led to a decrease of c-Fos protein abundance (Figure 5C). Taken together, these results showed that **JNJ-64619178** potently inhibited EWSR1-ATF1-mediated gene transcription in CCSST cells leading to impaired CCSST cell growth.

**Figure 5.**
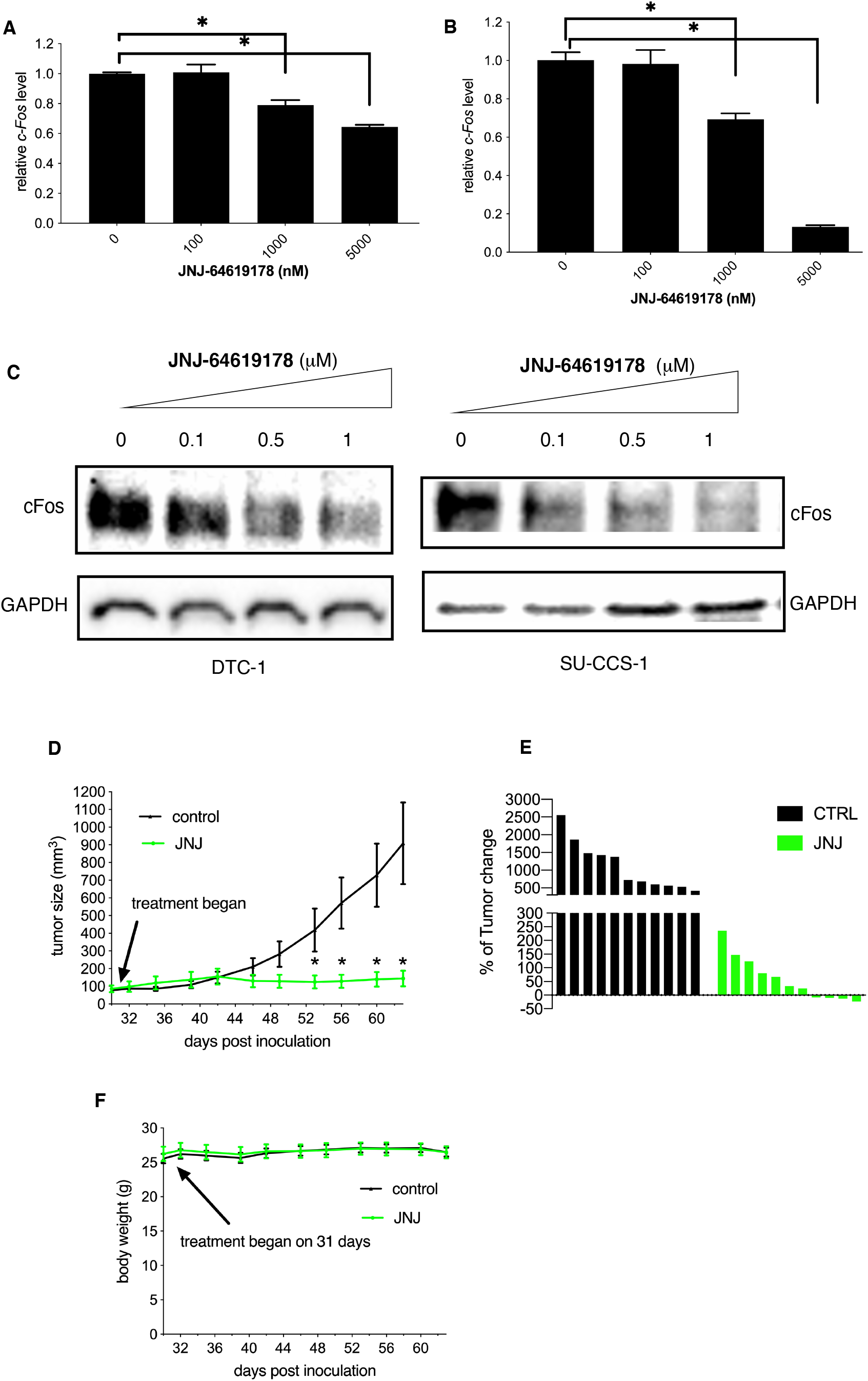
**JNJ-64619178** inhibited EWSR1-ATF1-mediated gene transcription and displayed anti-CCSST activity *in vitro* and *in vivo*. (A) **JNJ-64619178** inhibited *c-Fos* transcription in SU-CCS-1 cells. The cells were treated with different concentrations of **JNJ-64619178** for 24 h. Then the total RNA was isolated and converted into cDNA. The relative *c-Fos* transcript level was quantified by qRT-PCR using *18S* as a reference gene. (B) **JNJ-64619178** decreased *c-Fos* mRNA level in DTC-1 cells. The cells were treated with **JNJ-64619178** for 12 h. Then the total RNA was isolated and the relative *c-Fos* mRNA level was analyzed by qRT-PCR using *18S* as the reference gene. (C) **JNJ-64619178** decreased c-Fos protein level in SU-CCS-1 and DTC-1 cells. The cells were treated with different concentrations of **JNJ-64619178** for 24 h. Then the whole cell lysates were prepared for Western blot analyses using indicated antibodies. (D) **JNJ-64619178** efficaciously inhibited SU-CCS-1 tumor growth *in vivo*. SU-CCS-1 cells were implanted subcutaneously in athymic nude mice (male:female = 1:1). When the tumor sizes reached an average of ∼100 mm^3^, the mice were randomized to be treated with either vehicle or **JNJ-64619178** at 10 mg/kg (QDX5 weeks). The tumor volume was measured 2-3 times per week. (E) Waterfall plot of the tumor volume change at the end of treatment in (D). (F) **JNJ-64619178** treatment in mice did not change the body weights. The body weights of the treated mice in (D) were measured 2-3 times per week. * *P* < 0.05.

Encouraged by the promising activities seen with **JNJ-64619178** *in vitro* in inhibiting EWSR1-ATF1-mediated gene transcription and inhibiting CCSST cell growth, we further investigated its anti-cancer efficacy in an *in vivo* xenograft model. SU-CCS-1 cells were injected subcutaneously into immunodeficient mice (female:male=1:1). When the tumor was well-established and reached a size of ∼100 mm^3^, the mice were randomized to receive either vehicle or **JNJ-64619178** at 10 mg/kg by intraperitoneal injection. The daily treatment was lasted for 5 weeks and the tumor volumes were measured 2-3 times a week. As shown in Figure 5D, this tumor grew very aggressively when the mice were treated with a vehicle solution. On the other hand, when the mice were treated with **JNJ-64619178**, the tumor growth was essentially completely inhibited. While the tumors in the vehicle group grew 5-25-fold, the tumors in the treated group were either only slowly growing or regressed in this aggressive xenograft (Figure 5E).

During the entire course of treatment period, no change of body weight (Figure 5F) or other toxicity was observed, consistent with its safety profile observed in other models.^35^ These results support that the PRMT5 inhibitor **JNJ-64619178** represents a potential treatment option for CCSST.

## Discussion

CCSST remains a deadly disease with no cures. While the driver oncogene *EWSR1-ATF1* has been clearly identified, the disordered nature of EWSR1-ATF1 has made it recalcitrant for direct targeting using cell-permeable small molecules with appropriate drug-like properties. Due to its rare nature, it has been largely neglected to develop novel therapies. No standard systemic therapies exist. No targeted therapies have been identified as effective treatments for CCSST. No validated druggable targets have been identified for EWSR1-ATF1-driven CCSST. Previous studies have suggested that receptor tyrosine kinase c-Met might be a potential target for CCSST.^37^ However, c-Met has not been shown to modulate EWSR1-ATF1-driven gene transcription, the driver of CCSST.^37^ Furthermore, clinical studies using c-Met inhibitor crizotinib failed to produce meaningful benefits to CCS patients.^38^ Here, we identified PRMT5 as a critical protein to interact with EWSR1-ATF1 to enhance EWSR1-ATF1-mediated gene transcription and supporting CCSST cell growth, lending support that PRMT5 is a novel druggable target for EWSR1-ATF1-driven CCSST. Genetic silencing of *PRMT5* robustly inhibited CCSST tumor cell growth and EWSR1-ATF1-driven gene transcription. By targeting PRMT5 using different inhibitors, we further identified **JNJ-64619178** as a potential novel treatment option for CCSST. Interestingly, we found that the PRMT5 inhibitors based on **GSK591** are not effective in inhibiting EWSR1-ATF1-mediated gene transcription, suggesting that **JNJ-64619178** can uniquely impact the EWSR1-ATF1-PRMT5 complex at the gene promoter region to inhibit its transcription activity. It would be intriguing to investigate if the recently disclosed PRMT5 inhibitor PF-06939999 that is SAM-competitive^39^ has effect on EWSR1-ATF1-mediated gene transcription. Our results provide new insights into the role of PRMT5 as a transcription regulator. The different PRMT5 inhibitors with different modes of inhibition will be useful in further our understanding of PRMT5’s functions. Since the clinical development of **JNJ-64619178** for other oncology indications is ongoing (clinical trial #: NCT03573310), the results presented here provide compelling evidence to further clinically evaluate this PRMT5 inhibitor for EWSR1-ATF1-driven and deadly CCSST and other CCS-like malignancies.

### Materials and Methods

#### Plasmids and antibodies

The CREB1/ATF1 reporter construct CRE-RLuc was reported previously by us.^25^ Flag-tagged EWSR1-ATF1 expression vector was generated by In-Fusion^®^ Snap Assembly Master Mix (Takara) to fuse EWSR1 (2-325) in-frame with ATF1 (66-271) into pEGFP-N_3_ (Clontech). The EWSR1 (2-325) and ATF1 (66-271) fragments were PCR amplified from their respective open reading frames obtained from DNASU plasmid repository (Arizona State University). Lentiviral construct CRE-RLuc was prepared by gene synthesis followed by subcloning into pLJM1 vector (Addgene). All lentiviral shPRMT5 plasmids were purchased from Millipore Sigma. The lentiviral packaging vectors (pMD2.G and pMDLg/pRRE) were from Addgene. The following antibodies were used: anti-Flag (M2) (Sigma), anti-PRMT5 (Santa Cruz Biotechnology and Millipore), anti-GAPDH (Santa Cruz Biotechnology), anti-CREB1 (Cell Signaling Technology), anti-c-Fos (Cell Signaling Technology), anti-CBP (Santa Cruz Biotechnology).

#### Cell lines

HEK293T and SU-CCS-1 were obtained from ATCC (Manassas, Virginia). DTC-1 was a kind gift from Prof. Torsten Nielsen (University of British Columbia). CCS-292 was a generous gift from Prof. Charles Keller (Children’s Cancer Therapy Development Institute). The cells were maintained in Dulbecco’s Modified Eagle Medium (DMEM, Life Technologies) supplemented with 10% fetal bovine serum (Hyclone) and non-essential amino acids (Life Technologies). Mycoplasma contamination was tested regularly with MycoDect Mycoplasma Detection Kit (ALSTEM). All the cell lines have been authenticated using STR profiling.

#### Chemicals

**GSK591** and **GSK3326595** were obtained MedChemExpress LLC (Junction, NJ). **JNJ-64619178** was obtained from ChiralStar Inc (Parsippany, NJ). For *in vitro* assays, the drugs were dissolved in DMSO as stock solutions. For *in vivo* treatment, **JNJ-64619178** was dissolved in 1% *N*-methyl-2-pyrrolidone (NMP), 5% DMSO and 5% Tween-80 in ddI H_2_O.

#### Lentivirus preparation and transduction

The lentiviruses were prepared and used in a similar way as previously described.^40^ Briefly, lentiviruses expressing CRE-RLuc or indicated shRNA were prepared from HEK 293T cells by co-transfecting lentiviral expression plasmids along with packaging vectors using the calcium-phosphate method (TakaRa). DTC-1 cells were transduced with lentiviruses expressing CRE-RLuc. After puromycin selection, the cells were used directly for reporter assays.

#### shRNA silencing

SU-CCS-1 and DTC-1 cells were transduced with indicated lentiviruses expressing shCTRL or shPRMT5 for 24 h, when fresh media were added for another 48 h. Then the cells were used for the following experiments: cell growth assay by MTT, whole cell lysates for Western Blot and total RNA for qRT-PCR assays.

#### qRT-PCR

The cells were treated with indicated drugs. Then the cells were collected using the cell lysis buffer provided in NucleoSpin RNA (Takara). The total RNA was isolated following the manufacturer’s instructions. First strand cDNA was synthesized with PrimeScript 1st strand cDNA synthesis kit (Takara). Quantitative PCR reactions were performed using TB Green™ Advantage® qPCR Premix (Takara) in QuantStudio™ 7300 (Life Technologies). The 2^-Δ’Δ’Ct^ method was used to determine the relative expression level. The following primers were used: *18S*: 5′-GGATGTAAAGGATGGAAAATACA-3′ (forward), 5′-TCCAGGTCTTCACGGAGCTTGTT-3′ (reverse); *c-Fos*: 5′-GGAGGAGGGAGCTGACTGAT-3′ (forward), 5′-GAGCTGCCAGGATGAACTCT-3′ (reverse).

#### Transcription reporter assay

For transient transfections, HEK 293T cells were transfected with CRE-RLuc along with Flag-EWSR1-ATF1 using Lipofectamine2000^®^ (Life Technologies) following manufacturer’s instructions. Twenty-four hours post transfection, the cells were lysed using Renilla luciferase lysis buffer and the renilla luciferase activity was measured using Renilla luciferase assay system (Promega) in FB12 single tube luminometer (Berthold). For reporter assay in DTC-1 cells with stable CRE-RLuc integration, the cells were treated with indicated drugs at different concentrations for 24 h. The remaining renilla luciferase activity was measured as above. All the luciferase activity was normalized to protein concentration in each well and expressed as RLuc/μg protein.

#### IP-MS and IP-Western blot

HEK 293T cells were transfected with Flag-EWSR1-ATF1 using Lipofectamine2000^®^ (Life Technologies). Twenty-four hours post transfection, the cells were harvested and washed twice with cold PBS. Then the cell pellets were lysed in lysis buffer A (50 mM Tris, 5 mM EDTA, 150 mM NaCl, 1 mM DTT, 0.5% Nonidet P-40, pH 8.0) supplemented with protease inhibitor cocktail (Roche) and 1 mM PMSF. The cell lysates were pre-cleared with 1 μg mouse IgG (Rockland Immunochemicals) and protein A/G agarose beads (Pierce) for 1 h at 4 °C with tumbling. The pre-cleared lysates were then incubated with either mouse IgG or M2 for overnight at 4 °C. Then protein A/G agarose bead slurry was added to precipitate the immune complexes for 1 h at 4 °C. The bound immune complex was washed with lysis buffer A (3x) and the bound proteins were eluted from the beads using 1%SDS in PBS. The eluted proteins were analyzed by protein LC-MS/MS or Western blot.

#### Protein LC-MS/MS

Immunoprecipitated proteins from above were dried by vacuum centrifugation. They were then redissolved in 20 μL of 1x SDS-PAGE sample buffer, applied to wells of a NuPAGE 10% Bis-Tris SDS-PAGE gel (NP0301BOX), electrophoresed for 6 min at 200 V, and stained for 30 min with Imperial Blue protein stain (catalog no. 24615; Thermo Scientific). The gel was then rinsed in water and the entire top 1 cm of each lane containing proteins was excised, cut into 1 mm pieces, reduced/alkylated, and digested with trypsin for one hour at 50 °C in the presence of 0.01% ProteaseMax detergent using the method recommended from the manufacturer (Promega). Recovered peptides were then filtered using 0.22 µm Millipore Ultrafree-CL centrifugal filters. The filtrate was dried by vacuum centrifugation and then redissolved in 20 µl of 5% formic acid in preparation for mass spectrometric analysis.

Each digest was then chromatographically separated using a Dionex RSLC UHPLC system and delivered to a Q-Exactive HF mass spectrometer (Thermo Scientific) using electrospray ionization with a Nano Flex Ion Spray Source fitted with a 20 µm stainless steel nano-bore emitter spray tip and 1.0 kV source voltage. Xcalibur version 4.0 was used to control the system. Samples were applied at 10 µl/min to a Symmetry C18 trap cartridge (Waters) for 5 min, then switched onto a 75 µm x 250 mm NanoAcquity BEH 130 C18 column with 1.7 µm particles (Waters) using mobile phases water (A) and acetonitrile (B) containing 0.1% formic acid, 7.5-30% acetonitrile gradient over 60 min, and 300 nL/min flow rate. Survey mass spectra were acquired over m/z 375− 1400 at 120,000 resolution (m/z 200) and data-dependent acquisition selected the top 10 most abundant precursor ions for tandem mass spectrometry by HCD fragmentation using an isolation width of 1.2 m/z, normalized collision energy of 30, and a resolution of 30,000. Dynamic exclusion was set to auto, charge state for MS/MS +2 to +7, maximum ion time 100 ms, minimum AGC target of 3 × 106 in MS1 mode and 5 × 103 in MS2 mode.

Comet (v. 2016.01, rev. 2)^41^ was used to search 29,088 MS2 Spectra against a UniProt FASTA protein database (downloaded from www.uniprot.org November 2018) containing 21,080 canonical Homo sapiens sequences. An additional 179 common contaminant sequences were added, and sequence-reversed decoy entries of all sequences were concatenated to estimate error thresholds. The database processing used python scripts available at https://github.com/pwilmart/fasta_utilities.git and Comet results processing used the PAW pipeline^42^ from https://github.com/pwilmart/PAW_pipeline.git. Comet searches for all samples was performed with trypsin enzyme specificity with monoisotopic parent ion mass tolerance set to 1.25 Da and monoisotopic fragment ion mass tolerance of 1.0005 Da. A static modification of +57.02146 Da added to all cysteine residues and a variable modification of +15.9949 Da on methionine residues. Comet scores were used to compute linear discriminant function scores,^42, 43^ and discriminant score histograms were created for 2+, 3+, and 4+ peptide charge states. Separate histograms were created for target matches and for decoy matches for all peptides of seven amino acids or longer. The score distributions of decoy matches were used to estimate peptide false discovery rates (FDR) and set score thresholds at 2% FDR for each peptide class (10,335 peptide spectrum matches passed thresholds). Protein inference used basic parsimony principles and protein identifications required a minimum of 2 distinct peptide identifications per sample. Following filtering of common contaminants, this resulted in the identification of 540 proteins with one decoy protein match. The list of identified proteins is shown in Supplementary Table 1. The list of potential EWSR1-ATF1 interacting proteins was further filtered to consider only proteins having at least 3 unique peptides in the anti-Flag antibody group and 0 unique peptides in the IgG group, resulting in the identification of 6 proteins.

#### Western blot

The cells were treated with indicated drugs for different periods of time. Then the cells were harvested by scraping and washed with cold PBS twice. The cell pellets were lysed in lysis buffer A supplemented with 8 M urea. The protein concentration of the samples was determined using BCA assay kit (Pierce) and equal amount of proteins was loaded onto 4-20% SDS-PAGE gel (Bio-Rad) for Western blot analysis using indicated antibodies.

#### ChIP assay

The ChIP assay was performed using EZ ChIP Chromatin Immunoprecipitation kit (Millipore) following manufacturer’s instructions. Briefly, DTC-1 cells were crosslinked with 1% formaldehyde. Then the cells were lysed using provided SDS lysis buffer. The chromatin was sheared by sonication using FB120 probe sonicator (Fisher Scientific). The sheared chromatin was pre-cleared using mouse IgG and protein A/G bead slurry. The pre-cleared chromatin was then immunoprecipitated using IgG, anti-CREB1 or anti-PRMT5 along with protein A/G bead slurry. After extensive washes with Low Salt Immune Complex Wash Buffer, High Salt Immune Complex Wash Buffer, LiCl Immune Complex Wash Buffer and TE Buffer, the bound chromatin was eluted using 1% SDS in NaHCO_3_. The eluted chromatin was then reverse crosslinked and the resulting DNA was purified for qPCR analysis. The primers for *c-Fos* promoter were 5′-GGCCCACGAGACCTCTGAGACA-3′ (forward) and 5′-GCCTTGGCGCGTGTCCTAATCT-3′ (reverse).

#### MTT assay

The cells were plated in 96-well plates and were treated with indicated drugs at different concentrations for either 3 or 6 days. At the end of the treatment, the number of remaining viable cells was quantified using MTT reagent (Sigma, 0.5 mg/mL). The reduced purple formazan was dissolved in DMSO and absorbance values at 570 nm were obtained from an i3 multimode plate reader (Molecular Devices). The percent of growth is defined as 100 × (A_treated_ – A_initial_)/(A_control_ – A_initial_), where A_treated_ represents absorbance in wells treated with a compound, A_initial_ represents the absorbance at time 0, and A_control_ denotes media-treated cells.^36^ The GI_50_ was derived from non-linear regression analysis of the percent of growth–concentration curve in Prism 8.0.

### SU-CCS-1 xenograft assay

The use of animals was approved by OHSU Institutional Animal Care and Use Committee (IACUC). SU-CCS-1 cells were prepared as a 1:1 mixture in Hank’s Buffered Saline (HBS) and Matrigel (Corning). The 6-week-old athymic nude mice (Jax mice, female:male =1:1) were injected with 10 million cells subcutaneously to allow tumor intake. When the average tumor size reached ∼100 mm^3^, the mice were randomized to be treated with either vehicle or **JNJ-64619178** at 10 mg/kg once a day for 5 weeks (QDx5 weeks). The tumor volumes and body weights were measured 2-3 times per week. The tumors were measured in two dimensions using a digital caliper, and the tumor volume was expressed in mm^3^ using the formula V = 0.5*a*b^2^, where a and b represent the long and short diameters of the tumor, respectively.

### Statistical analysis

Student *t*-tests were used for calculating statistics, where a *P* value of less than 0.05 indicates significance. These tests were performed in Microsoft Excel (v 16.54).

## Acknowledgements

This work was made possible through financial supports provided from the National Institutes of Health R21CA220061 (XX), R01CA245964 (BL) and R01GM122820 (XX), Elsa U. Pardee Foundation, Oregon Health &Science University Technology Transfer Office, Oregon Health &Science University School of Medicine.

We thank OHSU Massive Parallel DNA Sequencing Core for authenticating the cell lines. The proteomic analysis was partially supported by NIH grants P30EY010572 (LL) and P30CA069533 (LL).

